# Counting sperm whales and visualising their dive profiles using two-hydrophone recordings and an automated click detector algorithm in a longline depredation context

**DOI:** 10.1101/2022.09.06.506716

**Authors:** Samuel Pinson, Gaëtan Richard

## Abstract

Odontocetes depredating fish caught on longlines is a serious socio-economic and conservation issue. A good understanding of the depredation behaviour by odontocetes is therefore required. Within this purpose, a method is described to follow diving behaviour of sperm whales, considered as proxy of their foraging activity. The study case took place around Kerguelen Islands from the Patagonian toothfish fishery. The method uses the coherence between direct-path sperm whale clicks, recorded by two synchronized hydrophones, to distinguish them from decoherent clicks that are reflected by the water surface or seefloor (due to surface roughness). Its low computational cost permits to process large dataset and bring new insights on sperm whales behaviour. Detection of sperm whale clicks permits to estimate the number of sperm whales and to assess their diving behaviour. Three diving behaviour were identified as “*Water Column*” (individual goes down and up), “*Water Wander*” (individual seems to go up and down multiple times in the water column), and “*Seafloor*” (individual spend time on the seabed). Results suggest that sperm whales have different diving behaviours with specific dives as they are either “*interacting*” or “*not-interacting*” with a hauling vessel.

## 1. Introduction

The intensification of fishing activity and rarefaction of resources has led to an increase in competition with marine predators worldwide over the last decades (Northridge and Hofman 1999; Read 2008). Such competition has led some marine predators to learn how to remove fish from lines or nets. This behaviour, named depredation, impacts substantially fishing activity, since it increases fishing costs and fishing efforts to finish the quotas (Gillman et al. 2006; Peterson et al. 2014; Richard et al. 2017) depredating species, with risk of lethal retaliation or entanglement (Donoghue et al. 2002; Gillman et al. 2006; Read 2008), and for the fish stocks, since depredation is not often accounted in fishery stock assessments (Peterson and Hanselman 2017). Although most fisheries are impacted by a broad range of large marine vertebrates depredating, most cases involve odontocetes interacting with longlines (Donoghue et al. 2002; Gillman et al. 2006; Northridge and Hofman 1999; Read 2008; Werner et al. 2015; Reeves et al. 2013). Longlines are composed of snoods (thin cords) connecting unprotected hooks along a mainline, increasing fishers’ selectivity by catching only targeted fish species, but at the same time making a resource easily accessible for marine predators (Gillman et al. 2006; Read 2008). Longlines could be deployed either within the water column, named pelagic longlines, or set on the seafloor, named demersal longlines. The fishing process is composed of three phases: (i) the setting, when hooks are baited and longlines are deployed at sea; (ii) the soaking, when the fish are caught as the longline is left at sea with no boat activity nearby; and (iii) the hauling phase, when the longline is recovered on board.

Depredation behaviour on longlines is highly variable depending on the type of gear (pelagic or demersal) and on the odontocete species involved. Pelagic longlines deployed close to the surface are always easily accessible to odontocetes and depredation appears to occur throughout the whole fishing process (Dalla Rosa and Secchi 2007; Forney et al. 2011; Passadore et al. 2015; Rabearisoa et al. 2012; Thode et al. 2016). On the other hand, demersal longlines are thought to be mostly depredated during hauling (Guinet et al. 2014; Hucke-Gaete et al. 2004; Mathias et al. 2012; Straley et al. 2014; Towers et al. 2019; Roche et al. 2007). However, recent studies suggested that sperm whales and killer whales may also depredate from demersal longlines during soaking (Richard et al. 2020, 2022; Janc et al. 2018; Cieslak et al. 2021; Towers et al. 2019). Although proofs that killer whales actually depredate on soaking demersal longlines are still not clearly confirmed, pieces of evidence for sperm whales doing so are clearer. Indeed, (Janc et al. 2018) revealed that an increasing soaking time was associated to higher numbers of sperm whales at hauling, suggesting that individuals arrived at the longlines during the soaking phase. Alternative types of data such as videos, passive acoustic monitoring and bio-logging are required to better understand the underwater dimension of depredation, notably during soaking. Additionally, (Towers et al. 2019) revealed positive correlations between the maximum dive depths of sperm whales, obtained from time-depth recorders, and the depths of nearest longline, suggesting potential depredation on soaking longlines too. Finally, (Richard et al. 2020) observed plausible evidence of sperm whale depredation on demersal soaking longlines using accelerometers fixed on longlines’ hooks, with an event confirmed by the entanglement of a sperm whale. Indeed, settings have specific acoustic signature which could attract attention of odontocetes before soaking (Richard et al. 2022). Nevertheless, this study was based on a limited dataset and the occurrence of depredation behaviour on soaking longlines could not be quantified.

In order to better describe and quantify depredation on soaking longlines, passive acoustic monitoring (PAM) is of great interest as it may cover a longer temporal scale around soaking longlines. Such approach was recently used to describe a common foraging activity by killer whales around soaking longlines (Richard et al. 2022). However, this study only relied on presence/absence of acoustic signals produced by killer whales, and there was no confirmed evidence of killer whales foraging on the seafloor or depredation on soaking longlines. Different setup in PAMs, using for instance array of several hydrophones, could nevertheless target this issue by method of spatial localisation of sound. Studies about the depredation issue in Alaska (Thode et al. 2015; Mathias et al. 2013) resulted in the description of sperm whales dive profiles around hauling longlines. Although sperm whale are likely to depredate on seafloor, how often remains unknown (Richard et al. 2020).

In this paper, the focus is on the sperm whale activity during the whole fishing process, i.e. from setting to soaking as in previous studies on killer whales (Cieslak et al. 2021; Richard et al. 2022). A passive acoustic monitoring method is described, allowing the dive behaviour of sperm whales to be visualized and classified, and thus to be used as a proxy of foraging activity. The objective is to correlate such behaviour to soaking demersal longlines, when no concurrent visual observations are available. The study case took place in the Kerguelen Economic Exclusive Zone (Southern Ocean) from the Patagonian toothfish fishery. The method uses the coherence between direct-path sperm whale clicks, recorded by two synchronized hydrophones, to distinguish them from decoherent clicks that are reflected by the water surface or the seafloor (due to surface roughness). The automatic detection of sperm whale clicks is thus based on recorded signals spatial coherence and has a low computational cost. However, the distance to and dive depth of the whales could not be determined due to uncertainties in the geometry of the two hydrophones.

The paper is organized as follows. The experiment and recordings are described in the section *Material* (2). The section *Methods* (3) describes the signal-processing method for automatic click detection (3.1) and how it can be used to analyze the data (3.2) which allows to interpret some diving behaviours (3.3). The *Results* section (4) focuses on the whole experiment campaign data analysis which is argued in the *Discussion* (5).

## 2. Material

The experiment took place in January 2017 within the economic exclusive zones of Kerguelen (49°0’S, 70°20’E) from a fishing vessel. Sound recordings and visuals observations were performed during fishing operations. Fishing regulations prohibit setting during daylight to avoid seabird bycatch (Weimerskirch et al. 2000). Several longlines were set at night and primarily hauled during the day. Longlines can stay from 6 hours to several days at sea. During longline hauling, trained fishery observers recorded interactions with sperm whales, defined as presence repeatedly diving whales within an approximate 500 m to 1 km range from the vessel (Roche et al. 2007; Tixier et al. 2010) (data available through the Pecheker database, accessible from the Natural History Museum of Paris (Martin and Pruvost 2007)). Longline positions (latitude and longitude), seafloor depths (500 – 2,500 m), setting times and hauling times were recorded.

An autonomous recorder (EA-SDA14, RTSys), paired with two hydrophones, was deployed on a longline. The acoustic recorder was clamped at 100 m deep on the downline connecting the buoy to the anchor (Fig. 1). The hydrophones were set 4.5 m apart vertically. However true geometry is unknown because of longline motions due to sea-surface waves and water currents. The recorder was programmed to record continuously during the whole longline deployments (i.e. from 6 hours to several days) with a sampling rate of 39 kHz. During the recording period, the fishing boat was moving independently of the longline provided with the acoustic recorder, and continued the fishing (i.e. setting and hauling other longlines).

**Figure 1.**
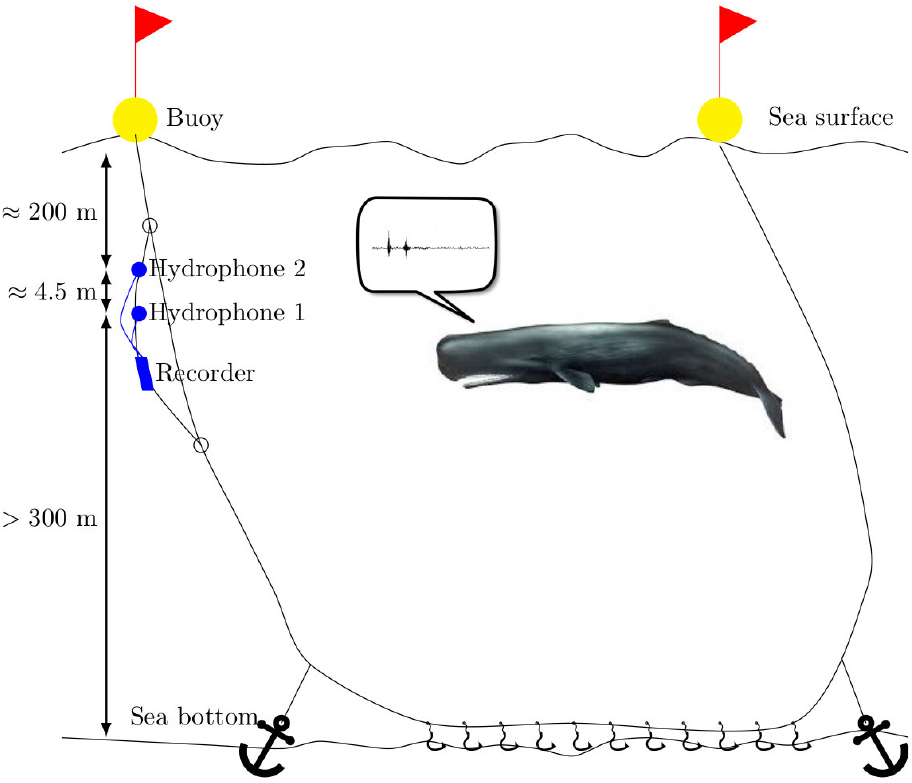
Sketch of a longline deployment on which 2 hydrophones and a recorder are attached.

## 3. Methods

The idea of the method is to take advantage of spatially incoherent reflections from the sea surface and the sea bottom (due to surface roughnesses) to identify the sperm whale click direct path in recorded signals. Indeed it is possible for reflected paths to be higher in amplitude than the direct one and amplitude-only click detectors may not be able to discriminate those different kinds of recorded clicks. Thus using two hydrophones which distance between them is sufficiently large, it is possible to identify the direct path from the reflected ones through their loss of spatial coherence. So it is hypothesized that the distance between the hydrophones is big enough compared with the dominant signal wavelength, to decorrelate sea-surface and sea-bottom reflections recorded by the two hydrophones.

Once the sperm whale click direct paths are detected, the goal is to image the successions of signal portions showing the time delay evolution of wave-guide reflections relative to the direct path.

### 3.1. Signal processing

Sperm whale clicks are described in details by Zimmer et al. (2005). They are generated by the phonic lips in the front of the forehead, that initially transmits the click backwards through the spermaceti organ, towards the frontal sac, which acts like a mirror and reflects the click forward through the “junk”. Thus sperm whale clicks are composed of pulses which result from multiple internal reflections within the sound generator in the forehead of the whale. So they are separated by a fixed interval that depends on the size of the forehead. Successive pulses (called P0, P1, P2, etc) have a frequency band between 3 kHz and 15 kHz. Additionally a low-frequency component below 3 kHz is present at P0. An example of a recorded click is presented in Fig. 2a with its time-frequency analysis in Fig. 2b where the two first pulses P0 and P1 are visible at *t* = 0 and *t* = 8 ms. The low frequency component associated with the P0 pulse is not visible in Fig. 2b because of the acoustic signature of the fishing vessel (i.e. the broadband noise below 3 kHz).

**Figure 2.**
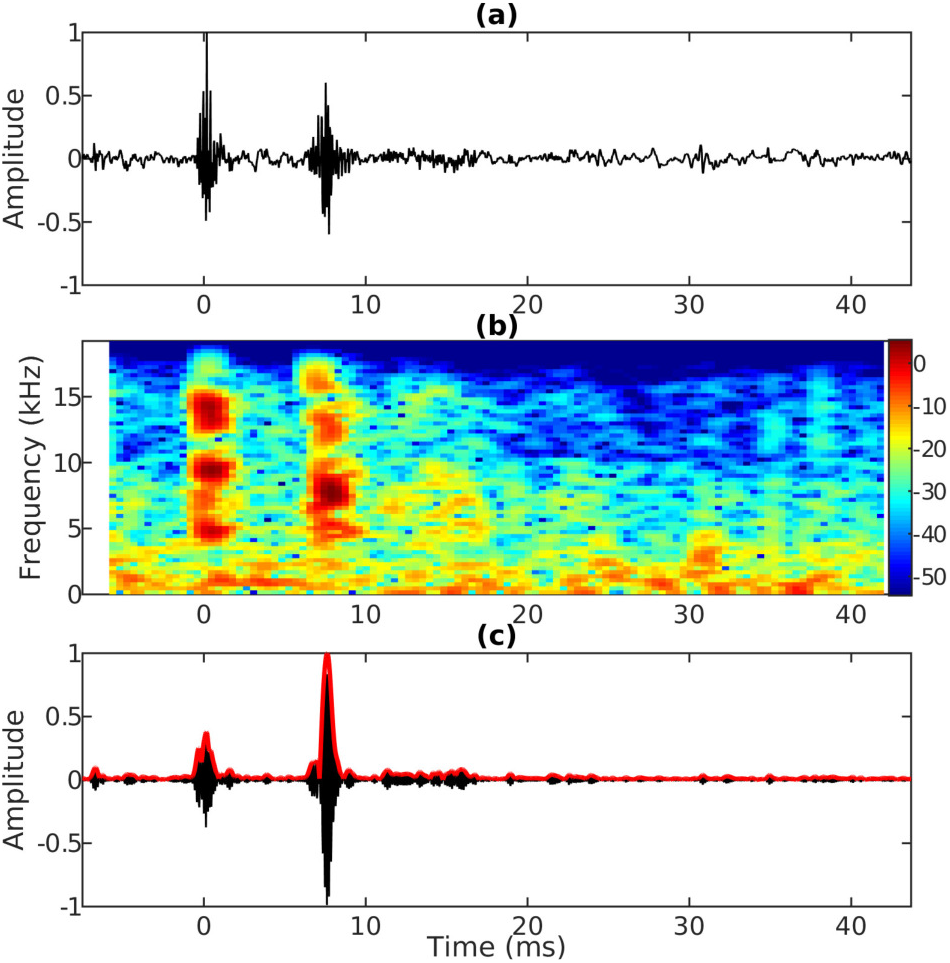
(a) Example of a recorded sperm whale click and (b) its time frequency analysis (color scale in dB with arbitrary reference). (c) Real part of the filtered signal (thin line) and its envelop (thick line).

For detection purposes, the recorded signals are modified into analytical signals. To do so the signal frequency spectrum that has to be narrow band, is filtered with a Gaussian function centered at *f* = 7.5 kHz and with 2.5 kHz frequency band (at −6 dB). This filtering also remove most of the boat noise energy below 3 kHz. Thus the recorded signals *p*_1,2_ from hydrophones 1 and 2 are preprocessed in the following manner:

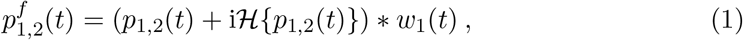

where * is the convolution operator, 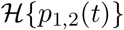 is the Hilbert transform of *p*_1,2_(*t*) and where the Fourier transform of *w_1_*(*t*) is the filter described above. Real part and absolute value of the filtered click is presented in Fig. 2c.

Coherence between two signals is usually an integral of the product of these two signals with a varying time delay between them and divided by the signal autocorrelations. But here the pattern to be detected is composed of two pulses P0 and P1, so the coherence between 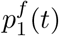 and 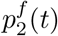 needs to be integrated only over the duration of a pulse by using a sliding window *w*_2_(*t*). This sliding time window is chosen to be a 4 ms long (at mid-amplitude) Gaussian function to be able to frame individual pulses. Thus the sliding coherence *Coh*(*t*, *τ*) used for click detection is:

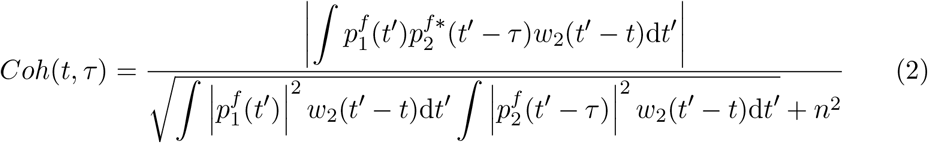

where 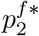 is the complex conjugate of 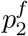, *τ* is the time delay between 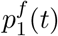 and 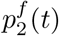 at which the coherence is evaluated, *t* is the time where the sliding window is centered on and, *n* is the signal noise level estimated from 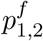 by fitting a Gaussian function with the signal amplitude histogram.

Fig. 3b shows the sliding coherence of the filtered signal in Fig. 3a where four clicks are visible and where the first one is the same as in Fig. 2a. Note that Equation 2 is evaluated only for |*τ*| < *d*_12_/*c* ≈ 3.2 ms where *d*_12_ ≤ 4.6 m is the distance between the hydrophones and *c* ≈ 1450 m/s is the water sound speed. One can distinguish echoes (likely from the sea-surface) about 180 ms after the direct paths in Fig. 3a. The sliding coherence shows in Fig.3b values close to one at direct path clicks. The sea-surface reflections are not detected because they are incoherent. It appears that there is some coherence for *τ* ≈ −1 ms. It is believed that this coherence comes from the fishing-boat cavitation bubble implosions for which the sounds are also coherent between the two hydrophones. As sperm whale clicks are also detected when emitted near the sea surface (see next section), it is hypothesized that Loyd mirror effect does not affect significantly the coherence. Indeed, the Fresnel zone size is getting smaller when the source (or receiver) is getting closer to the surface and thus, coherence may be less affected by the surface roughness.

**Figure 3.**
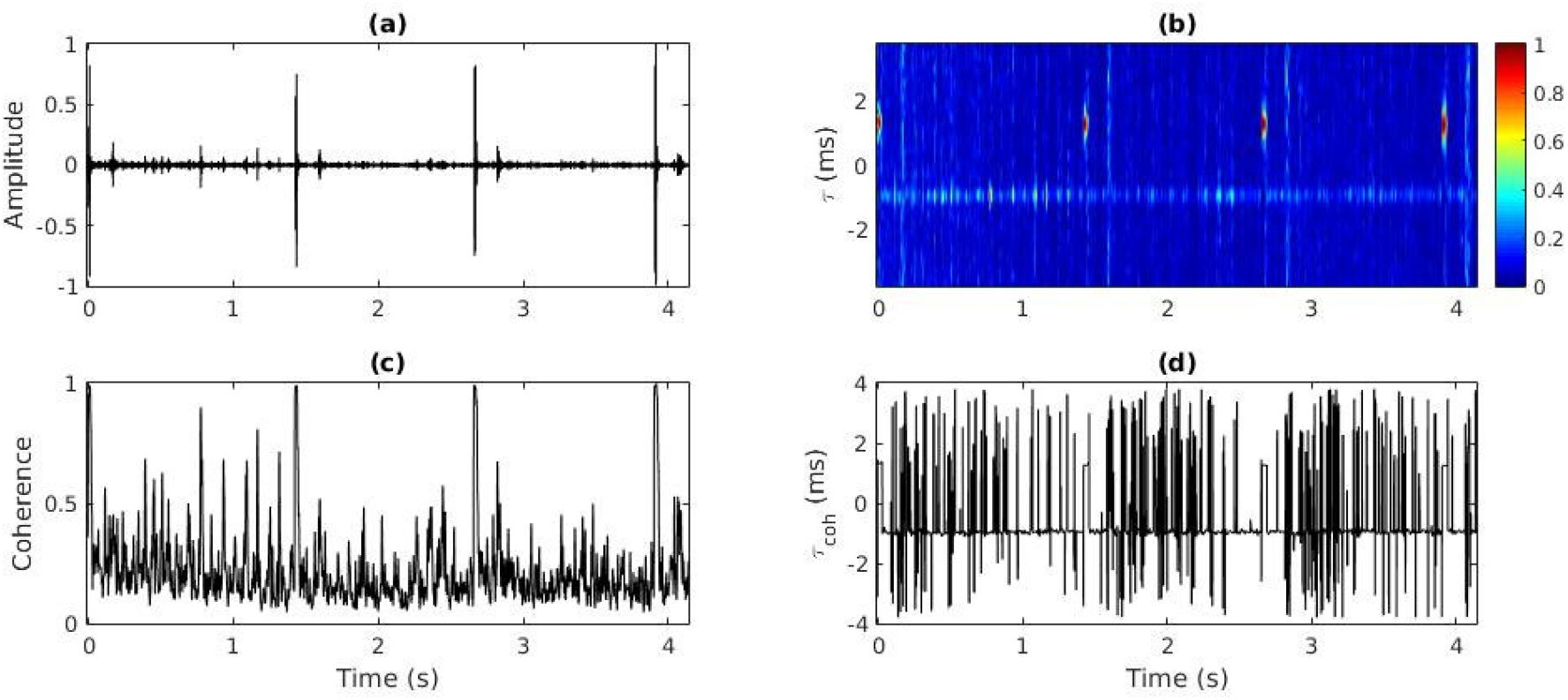
(a) Real part of the filtered signal 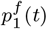 in which four sperm whale clicks associated with their sea-surface reflections (≈180 ms later) are visible and (b) its corresponding sliding coherence *Coh*(*t*, *τ*) in which clicks appear with a coherence close to one while sea-surface reflections are invisible. (c) The coherence signal *coh*(*t*) used for click detection and (d) the associated time delays between the two hydrophones *τ_coh_*(*t*).

Finally, the coherence to be used for click detection is:

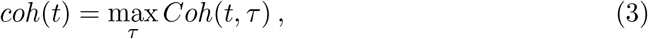

where the time delay for which the coherence is maximum is:

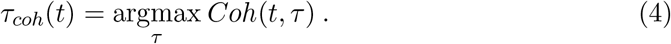

Fig. 3c shows the results for *coh*(*t*). Direct path clicks from the example show a coherence close to one (the other spikes are believed to originate from fishing boat cavitation noise). Fig. 3d shows *τ_coh_*(*t*) for which the coherence is maximum. It appears somehow random but its value only has meaning when *coh*(*t*) is close to one.

The click arrival time is defined at the maximum of amplitude of the P0 pulse. The coherence function *coh*(*t*) creates a window in which looking for P0 and P1 pulses. Nevertheless, looking for two local maxima corresponding to P0 and P1 in 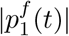 (thin line in Fig. 4) is difficult because it presents rapid fluctuations within each pulse. So 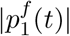 is convolved with a smoothing Gaussian window:

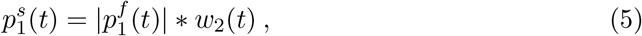

where *w*_2_(*t*) is the same window as for the sliding coherence. Thus 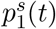 (thick line in Fig. 4) has its two maximum amplitude peaks corresponding respectively to P0 and P1. One can see that both pulses P0 and P1 are contained by the coherence (dashed line in Fig. 4 which corresponds to the coherence in Fig. 3c). The first step of the algorithm to detect sperm whale direct path clicks is to fix a coherence threshold (fixed at 0.6 empirically) to frame signal portions that will be identified as coherent. Fig. 3c shows that some boat-noise spikes are higher than 0.6. It appeared that those spikes are shorter than sperm whale clicks. So the second step is to remove all coherent portions shorter than 10 ms (fixed empirically). This second step is sufficient to remove most of the coherent portions due to the fishing boat. The third step is to locate the two highest local maxima within each coherent portion of 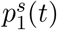. The first local maximum is then identified as the P0’s arrival time. Indeed the relative amplitude between P0 and P1 depends on the whale orientation (Zimmer et al. 2005) and thus the maximum may be either P0’s or P1’s.

**Figure 4.**
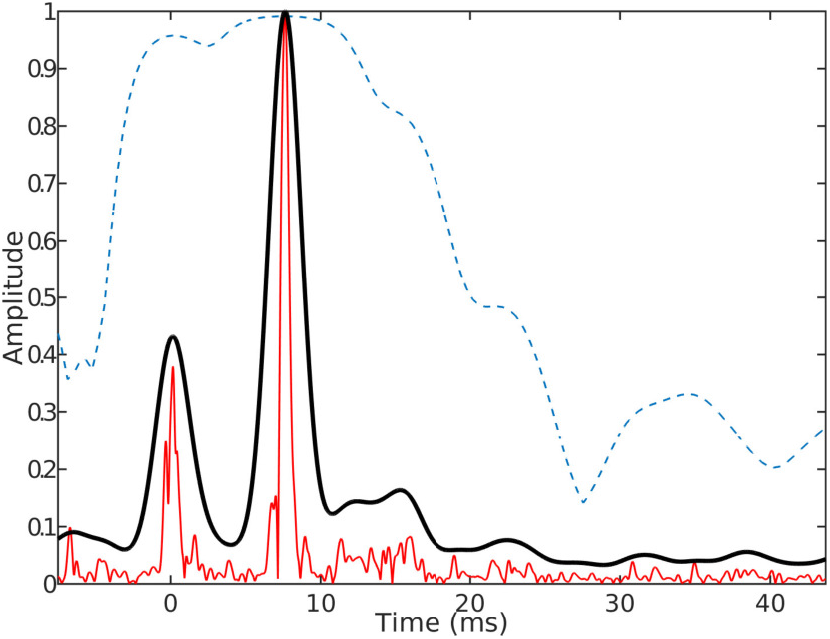
(thin line) The filtered signal envelop 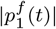 of the sperm whale click presented in Fig. 2a and (thick line) its smoothed version 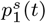. The dashed line is the coherence signal *coh*(*t*).

### 3.2. Data analysis

Once each direct path click arrival time is identified by the detection algorithm, it is possible to store short portions of 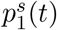 for which *t* = 0 correspond to the P0 local maxima and display the relative time of arrival of the boundary reflections. The result obtained from the 01/20/2017 18-hour record is presented in Fig. 5b and the corresponding *τ_coh_* are plotted in Fig. 5a. In principle, it should be possible to calculate dive depth of and distance to the whales from Fig. 5b, by analyzing relative travel times of sea surface and seafloor echoes (Thode 2004, 2005) but the uncertainties on the hydrophone positions here are too important to make it possible. Nevertheless it has been possible to find three different sperm whale dive patterns in Fig. 5b. The first one (Fig. 5c) is a simple dive into the water column and corresponds to a monotonic increase of sea-surface echo travel time followed by its monotonic decrease, named “*water-column dive*”. The second pattern (Fig. 5d) named “*seafloor dive*” has a similar sea-surface echo evolution, plus a sea-bottom echo crossing it (at 1:08 in Fig. 5d) and then merging with the direct path (at 1:18 in Fig. 5d) meaning that the whale dove down to the sea-bottom. The third pattern (Fig. 5e) suggests that the whale is wandering in the water column as the sea-surface echo evolves back and forth, and thus named “*wander dive*”. Dive behaviour is sometimes hard to identify in Fig. 5b as some echoes from different whales are entangled. Indeed, it is possible to identify numerous whales in Fig. 5a as direct-path time delays follow individual trajectories. In particular, one can identify 7 different trajectories between 11:00 and 13:30 where the individuals seem to split up before gathering again.

**Figure 5.**
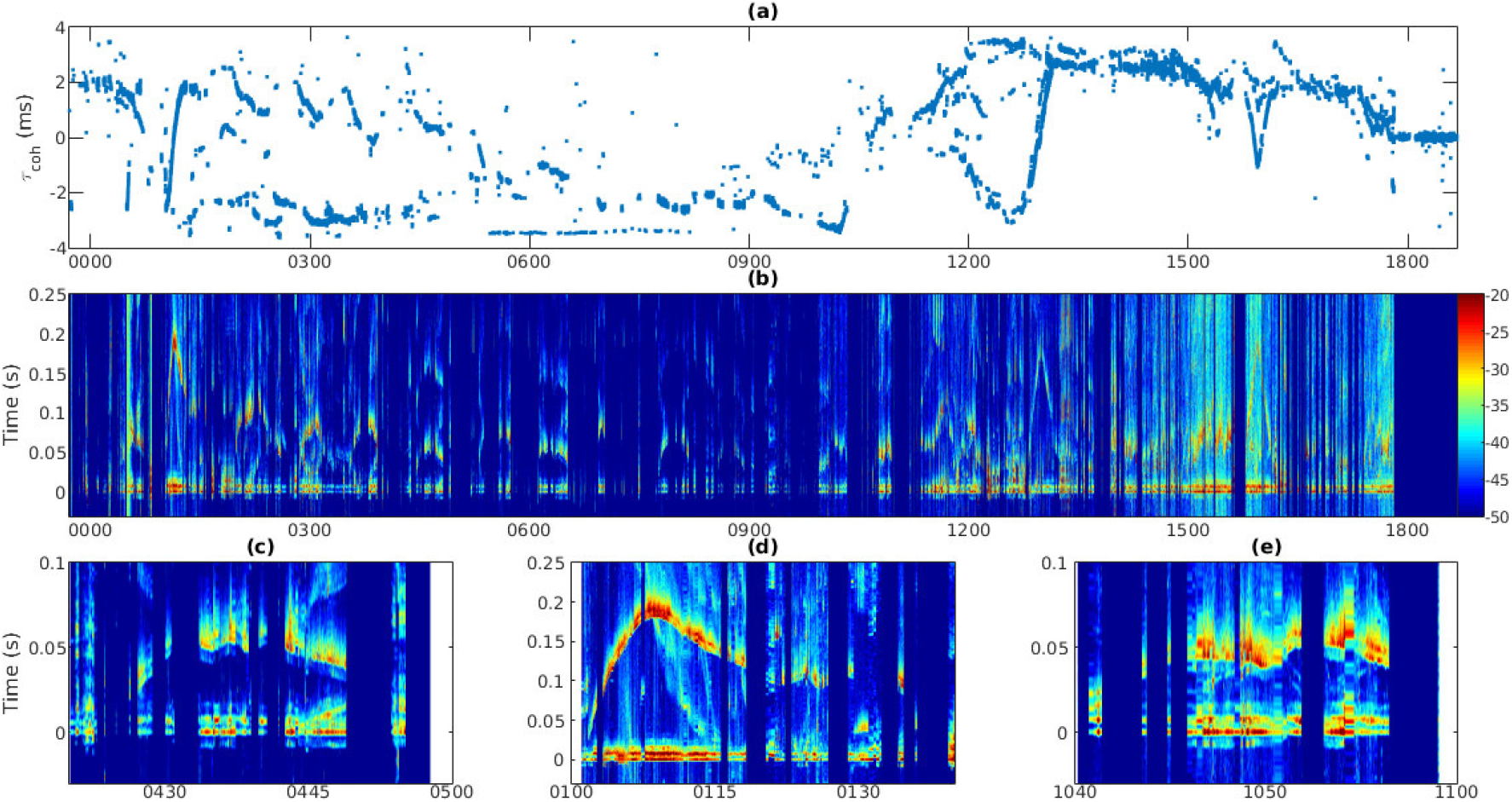
(a) Click time delays *τ_coh_* between the two hydrophones of the detected clicks. Numerous whales can be identified by the time delays following individual trajectories, e.g. between 11:00 and 13:30, where 7 different trajectories indicate that the individuals seem to split up before convening again. (b) Short portions of 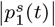 (colour scale in dB with arbitrary reference) for which the vertical axis is the time relative to the detected direct-path clicks. Signal portions are displayed horizontally as a function of the acquisition time. (c) *Water column dive*, where the surface echo delay times monotonically increase while the whale dives, and then smoothly decrease when the whale ascends. (d) *Sea floor dive*, where the surface echo delay times increase during descent to 200ms at 1:08. At this point weaker sea bottom echoes, starting with a 200ms delay appear, then smoothly decrease to zero at 1:18 when the whale reaches the bottom. During this time interval, the sea surface reflections smoothly decrease to 125ms probably due to outgoing whale displacement. (e) *Wander dive*, where the surface echo time delays fluctuate while the whale changes dive depth.

### 3.3. Diving behaviour analysis

Using a descriptive approach, diving behaviours described in previous section 3.2 are associated with fishing activities (Fig. 6a). The fishing vessel can be either setting, hauling or travelling (i.e., not engaged in any fishing activities). Visually observed and acoustically detected sperm whales are considered to be the same individuals if the fishing vessel (where observers are) is within the acoustic detection range, of about 30 km (Mathias et al. 2013), from the longline equipped with the hydrophones. These identified dives were then considered as “*interacting with hauling*”. Conversely, dives were considered as “*not interacting with hauling*“, if at least one of the three following conditions is met: (*i*) the fishing vessel was either setting or travelling; or (*ii*) no whale was visually detected from the fishing vessels; or (*iii*) the visual observation occurred outside the acoustic detection range of sperm whale clicks from the hydrophone. Although acoustically detected whales could not be positively identified as the one visually observed from the fishing vessel, individuals observed from vessels within the acoustic detection range of clicks were likely to be recorded by the hydrophones. Nevertheless, dives not associated with a hauling vessel (i.e. *not interacting with hauling*) were more confident since no whale was observed from the fishing vessels or within the acoustic detection range of clicks.

**Figure 6.**
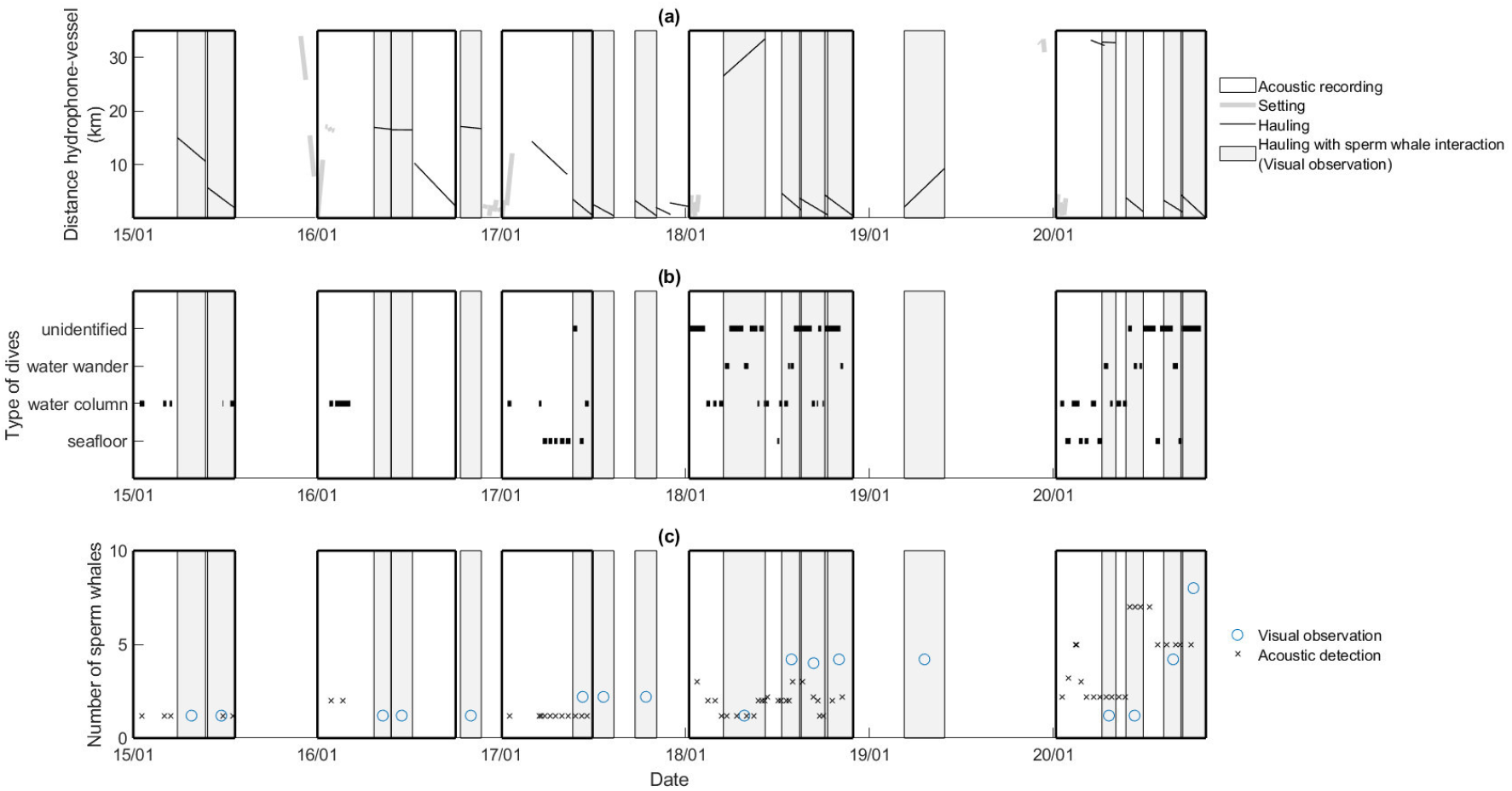
(a) Distance between recorder and fishing activities during the study period (January 15-20). Thick gray lines correspond to setting of long lines and thin black lines to hauling. Boxes with thick lines mark sound recordings, boxes with grey shade indicate visual observations of sperm whale interactions during hauling. (b) Diving behaviours (wander, column and seafloor dives) as identified from sound recordings. (c) Number of visually (open circle) and acoustically (x) identified whales during the study period.

## 4. Results

### 4.1. Sperm whale counting and behaviour analysis

A total of 81.6 hours of acoustic data was recorded in connection with five deployments (mean duration of 16.3 ±4.1 hours per deployment). In these recordings a total of 51 dives were identified, with a total duration of 25 hours (mean duration of 0.5 ±0.3 hours per dive). In 12 events, with a total duration of 17 hours (i.e. ≈ 40 % of the total dive; mean duration per event 1.4 ± 0.8 hours) the dive types could not be identified due to clicks from different whales being superposed, making it impossible to associate the correct surface or seafloor reflections. Among the 51 identified dives, 57 % were associated as water-column diving behaviour (Fig. 5c and Fig. 6b), 25 % as seafloor diving behaviour (Fig. 5d and Fig. 6b) and 16 % as wander-dive behaviour (Fig. 5e and Fig. 6b).

The number of sperm whales for every observation varied from 1 to 7 individuals (Fig. 6c). Counting from acoustic detection was similar to the number of individuals estimated by observers from the fishing vessel at hauling (Fig. 6c). However, the last recording revealed that during three haulings the acoustic method estimated more individuals than the visual observation from the fishing vessel (Fig. 6c). Additionally, the acoustic method also allows for estimating the number of individuals in the area, within detection range of ca. 30km, when no visual observations are carried out, e.g. during setting and travel.

### 4.2. Sperm whale interaction with boat

More diving sperm whales were detected as *not interacting with hauling*, n=30, than as *interacting with hauling,* n=21 (Fig. 7a). When *not interacting with hauling,* the whales seemed mainly engaged in *water column dives,* which constituted 63 % of the totally 30 dives. The other 37 % dives were to the seafloor (Fig. 7a), since no wander behaviour was detected at these time (i.e. *not interacting with hauling*). Besides, the majority of *seafloor dives,* 11 dives representing 85 % of all such dives, were associated with *not interacting with hauling*. Seafloor dives were then almost exclusive to *not interacting with hauling* behaviour (Fig. 7b). Conversely, wander dives occurred exclusively when *interacting with hauling* (Fig. 7b), and represented 43 % of the interacting dives (Fig. 7a). When *interacting with hauling,* the whales were mainly engaged in *water column dives,* representing 48 % of all 21 dives. Only two *seafloor dives* (i.e. 15 % of these dives) were identified in connection with *interacting with hauling* (Fig. 7a). Altogether, these results suggest that sperm whales have different diving behaviours with specific dives as they are either *interacting with hauling* or *not-interacting with hauling*.

**Figure 7.**
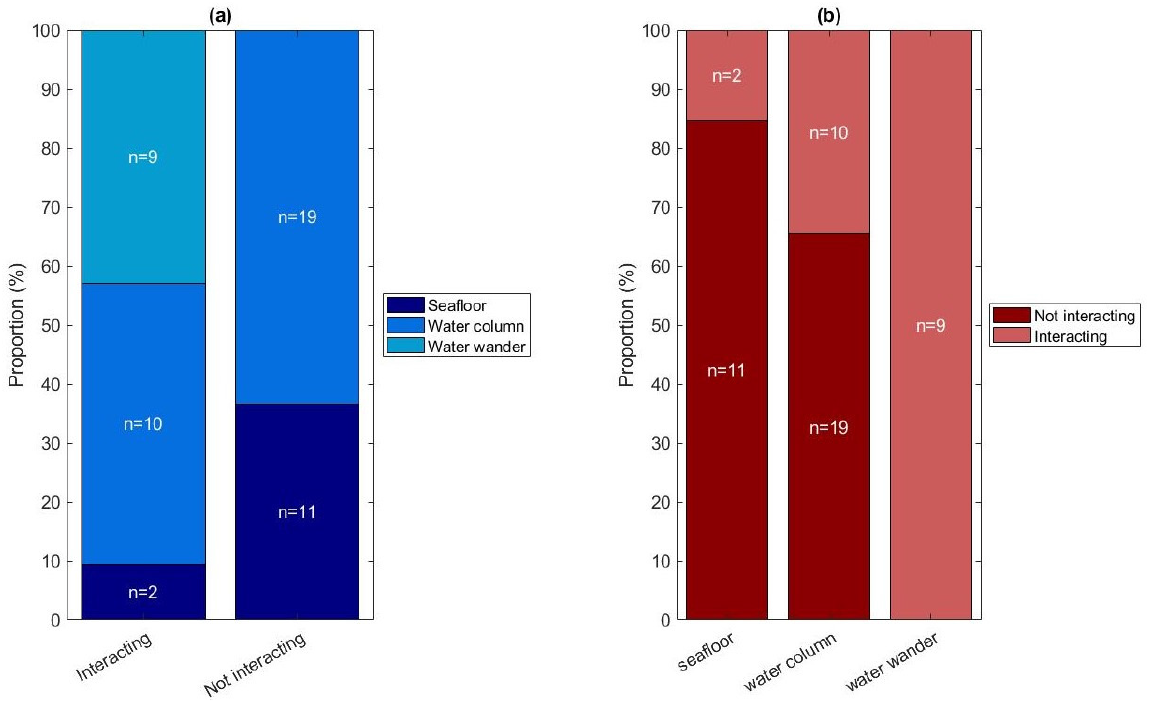
(a) Proportion of dive types in sperm whales interacting and not interacting with hauling. The majority of dives were *water column dives* (N=29), followed by *seafloor dives* (N=13) and *wander dives* (N=9). (b) Proportion of interacting vs not interacting with hauling in connection with the three dive types. Although most of the *water column dives* were not associated with hauling, an equal number of such dives and *wander dives* were associated with hauling.

## 5. Discussion

Taking advantage of the bad weather conditions by using a coherence function to distinguish the acoustic direct path from the reflected ones permits to automatically detect sperm whale clicks. As the technique is based on simple signal processing, the computation time is of about a quarter of the recording time on a laptop computer. In principle, it should be possible to estimate the range of the whale from the two hydrophones by using the angle of arrival of the direct click and the time delay of the sea-surface reflection (Thode 2004, 2005; Mathias et al. 2013). To do so, it is necessary that the hydrophones remains on vertical position and the distance between them stays steady. But such trivial theoretical requirements remain challenging in the real world, especially when deployments are logistically constraint by fishing operation in rough conditions as encountered in the Southern Ocean.

The passive acoustic method shows great potential to estimate the number of sperm whales around longlines, at least within a 30 km range from the two element hydrophone array. This acoustic approach could then be complementary to traditional visual counting from the vessel at hauling (Roche et al. 2007; Tixier et al. 2010). However, the acoustic method has the added advantage that whales can be counted also when the fishing vessels are not hauling, and thus have no visual observers in action (Richard et al. 2022). Also, in some of the deployments reported here, the acoustic method counted more whales than reported by the visual observers, indicating that the latter may underestimate the real hauling interaction rates. Indeed, missing the presence of some individuals is likely to understate interaction rates (Roche et al. 2007; Tixier et al. 2010). However, assessing whether depredation events are missed by visual observations requires a good monitoring of sperm whales’ behaviours around soaking longlines.

The second advantage with this passive acoustic method is that it allows for the identification of different dive behaviours. Although it turned out to be difficult to identify the different dive types when several whales are clicking simultaneously, still a large proportion of the whales in this study could be individually distinguished to allow for the identification of their dive types. The association of *wander dives* with hauling suggests that this may be a depredation-specific behaviour, where the repetitive descents and ascents could be the whale moving along the longline and raking it, as described in Alaskan fisheries (Mathias et al. 2009). Also, the higher proportion of *water column dives* than *seafloor dives* in connection with hauling is consistent with the shallow dive behaviour described for sperm whales depredating on Alaskan longline fisheries (Mathias et al. 2013). However, this dive type may also be a natural foraging behaviour for pelagic cephalopods (Whitehead 2009), since it was the main dive type in whales not interacting with hauling. In this case, foraging on pelagic prey would represent here 63 % of their diving behaviours when not interacting with a vessel, against 37 % of their dives considered as feeding on a demersal fish as Patagonian toothfish (Collins et al. 2010). Conversely, the higher proportion of seafloor dives when not interacting with a hauling vessel may either reveal a natural foraging behaviour on demersal prey as Patagonian toothfish (Abe and Iwami 1989) or a seafloor depredation behaviour (Richard et al. 2020).

Although the method does not enable to distinguish between natural foraging at seafloor or depredation behaviour on soaking longlines, the optimal foraging theory (Charnov 1976; Pyke 1984) would suggest that individuals are more likely to use the easiest resource, and thus that they may depredate on soaking longlines. Within this assumption, 22 %of the detected dives (i.e. 11 *seafloor dive* while *not interacting with hauling* among the 51 dives) in this study would then be associated as missed soaking depredation events (Richard et al. 2020). Conversely, 41 % of all dives were clearly associated to interaction behaviour at hauling. This observation may have major implications for estimation of interaction rates and perhaps even depredation rates. The real depredation rate is the difference in catch per unit effort, and may be the sum of both seafloor and hauling depredation (Tixier et al. 2010; Gasco et al. 2015). Hence, only counting possible hauling depredation may underestimate the depredation rate.

## 6. Conclusion

This passive acoustics method, based on two hydrophones and the application of coherence-based analysis, allows for the estimation of the number of sperm whales within a area of 30 km and the classification of their dive behaviour. Three dive types were identified: (i) *Water column dive,* where the whale monotonically descended in mid water, and then monotonically ascended, (*ii*) *seafloor dive* where the whales dove all the way down to the seafloor, and (*iii*) *wander dive* where the whale did multiple descent and ascents in mid water. These behaviours could be interesting proxies for a better understanding of interactions with fisheries. Future research should focus on two aspects. First a greater robustness of the acoustic recorder geometry would make it possible to estimate the depth and distance of the sperm whales. This would make it possible to better discriminate possible depredation from natural foraging by localising more precisely clicking individuals among the longlines and within the water column. Second, to increase the acoustic dataset as well as the visual observations, in order to refine the estimation of the interaction and/or depredation rates.

## Acknowledgments

We warmly thank the captains, their crew and the fishery observers for their help in the data collection. We also thank the Natural History Museum (Musée National d’Histoire Naturelle) of Paris for providing access to the PECHEKER Database. We are also very grateful for the support of the Fondation d’Entreprise des Mers Australes, the Syndicat des Armements Réunionais des Palangriers Congélateurs, fishing companies, the Direction des Pêches Maritimes et de l’Aquaculture, Terres Australes et Antarctiques Françaises (the Natural Reserve and Fishery units). We thank Dr. Julien Bonnel, Dr. Christophe Guinet and Dr. Flore Samaran for their help and contribution within the Orcadepred project. We warmly thank the anonymous reviewer from a previous submission who greatly helped improving the manuscript with his insightful comments.

## Ethical statement

All instrument deployments followed the ethics policies of the French Southern and Antarctic Lands (TAAF) and were authorized by the Réserve Naturelle Nationale (RNN des TAAF) through approval A-2017-154.

